# Artemisinin mimics nitric oxide to reduce adipose weight by targeting mitochondrial complexes

**DOI:** 10.1101/157396

**Authors:** Qian Gao, Jiang He, Tao Liao, Yan-Ping Chen, Li-Li Tan, Ji-Da Zhang, Chang-Qing Li, Qing Zeng, Qi Wang, Shui-Qing Huang, Xin-An Huang, Qin Xu, Qing-Ping Zeng

## Abstract

It remains obscure how to medically manage visceral obesity that predisposes metabolic disorders. Here, we show for the first time that a trace amount of artemisinin (0.25 mg/kg) reduces adipose weight in an inflammatory obese mouse model induced by a high-fat diet with lipopolysaccharide (HFD+LPS). HFD+LPS trigger pro-inflammatory responses, upregulate *NOS2* expression, elicit potent nitric oxide (NO) burst, and reinforce adipose mitochondrial dysfunctions that facilitate adipogenesis for visceral weight gain. By targeting mitochondrial complexes, artemisinin resembles the NO donor nitroglycerin to exert anti-inflammatory effects, downregulate *NOS2* expression, maintain stable NO release, and augment adipose mitochondrial functions that necessitate adipolysis for visceral weight loss. Taken together, artemisinin plays adipose weight-reducing roles by rectifying inflammation-driven mitochondrial dysfunctions.

---

The high-fat and low-fibre diets render overweight and obesity overwhelming throughout the world^1^. Even though weight reduction is often a hot-spot topic, it remains a dilemma in dealing with obesity and solving obesity-related health issues, such as noninsulin-dependent diabetes mellitus (NIDDM) and nonalcoholic fatty liver disease (NAFLD)^2^,^3^. The American Medical Association (AMA) has declared obesity a disease^4^, so it should be reasonable to clinically manage weight reduction. In this regard, the commonly used anti-diabetic drug metformin represents a favorable weight-loss medication^5^. Potentially, a slow released and low toxified 2,4-dinitrophenol (DNP) seems to hold promise in weight reduction by medication^6^. Mechanically, DNP allows protons to leak across mitochondria to uncouple oxidation from phosphorylation, suggesting mitochondrial uncoupling might be a mechanistic target for treatment of obesity^7^. Metformin dually activates adenosine monophosphate (AMP)-activated protein kinase (AMPK)^8^ and oxidized nicotinamide adenine dinucleotide (NAD^+^)-dependent silent mating type information regulation 2 homolog 1 (SIRT1)^9^, both in turn coordinately activate peroxisome proliferators-activated receptor gamma coactivator 1-alpha (PGC-1α) that necessitates mitochondrial biogenesis^10^. It is therefore certain that DNP and metformin exert weight-reducing effects by impacting on mitochondria and modulating their energetic functions.

It was suggested to categorise obesity to subcutaneous and visceral adipose depot because the former is metabolically “healthy” with the normal scores on the indices of insulin resistance and systemic inflammation, whereas the latter is metabolically “unhealthy” in an association with metabolic diseases^11^. However, it remains largely unknown whether both types of obesity are equally induced once nutrient intake is more than energy expenditure. To answer this question, we suggest here a hypothesis of “pro-inflammation/anti-inflammation-switched high/low-level nitric oxide (NO) to modulate visceral adipogenesis/adipolysis”. The pro-inflammatory signals were suggested to upregulate *NOS2*/inducible nitric oxide synthase (iNOS) expression, trigger potent NO burst, permanently block mitochondrial respiration, and augments energy deposition. In contrast, the anti-inflammatory responses were assumed to downregulate *NOS2*/iNOS expression, maintain mild NO release, frequently enhance mitochondrial biogenesis, and promote energy expenditure. As supporting evidence, it was shown that high-level NO inhibits cell respiration by binding to cytochrome *c* oxidase (COX), whereas slow and small-scaled NO release stimulates mitochondrial biogenesis in diverse cell types^12^. We also found the pro-inflammatory cytokines, tumor necrosis factor α (TNF-α) and interleukin 1β (IL-1β), upregulate iNOS expression and potentate NO burst^13^,^14^.

AMPK can be activated even under an inflammatory condition because IL-1β treatment leads to increased iNOS expression, which occur concurrently with activation of AMPK^15^. It was recently demonstrated that AMPK catalyses the phosphorylation of Janus kinase 2 (JAK2) and inhibit the pro-inflammatory JAK2-signal transducer and activator of transcription 3 (STAT3) signaling^16^, addressing metformin that activates AMPK might also exert an anti-inflammatory effect. Indeed, metformin was proven to impede the pro-inflammatory STAT3 and NF-κB signaling^17^. In similar, calorie restriction (CR) that activates AMPK and mitigates inflammatory responses was validated to foster weight loss and benefit lifespan extension^18^. Besides, AMPK was also unraveled to activate *NOS3*/endothelial nitric oxide synthase (eNOS) via an AMPK→Rac1→Akt→eNOS pathway^19^. Therefore, it would be said that AMPK can improve mitochondrial functions by downregulating iNOS and activating eNOS, and AMPK activators that target mitochondria and thus elevate AMP levels are, or can be developed to, weight-reducing drugs once their doses are optimally chosen.

Artemisinin, an antimalarial sesquiterpenoid with a moiety of endoperoxide lactone derived from the medicinal herb *Artemisia annual* L., was found to target yeast mitochondrial NADH dehydrogenase, whose deletion reduces yeast’s sensitivity to artemisinin^20^. Artemisinin was also revealed to target malarial mitochondria for eliciting generation of reactive oxygen species (ROS), during which NADH dehydrogenase inhibitors behave as antagonists against artemisinin-induced ROS generation^21^. Additionally, artemisinin was identified to induce seven mitochondrial proteins in hepatocarcinoma cells^22^, and artemisinin-bound malarial mitochondrial NADH dehydrogenase was formulated by a theoretical model^23^. In our previous work, conjugation of artemisinin with haem and synchronous induction of AMPK and COX were monitored in yeast and mice, suggesting artemisinin triggers mitochondrial biogenesis by conjugating COX and activating AMPK^24,25^.

To experimentally validate whether artemisinin would possess a potential weight-reducing role, we applied artesunate (AS), a semi-synthetic derivative of artemisinin, to explore a possibility of adipose weight reduction in a mouse obese model induced by a high-fat diet (HFD) combined with lipopolysaccharide (LPS). AS was chosen because it was known to mimic NO to initiate mitochondrial biogenesis^24,25^. By localizing AS binding to the heme protein cytochrome c1 (CYC1) of the mitochondrial complex 3 and the non-heme protein NADH dehydrogenase ubiquinone flavoprotein 1 (NDUFV1) of the mitochondrial complex 1, we disclosed for the first time that AS resembles the NO donor nitroglycerin (NG) to exert adipose weight-reducing effects. These encouraging results should shed light on the development of artemisinins as an alternative weigh-reducing drug based on the thorough elucidation of an insidious role of pro-inflammatory adipogenesis and a mechanistic link to anti-inflammatory adipolysis in the pathogenesis of metabolic syndromes.

## Results

### AS/NG reduces adipose weight with decrease in insulin and increase in leptin

To testify whether AS/NG would exert the effect on visceral weight reduction, we established an inflammatory obese mouse model by daily feeding HFD and intraperitoneally injecting 0.25 mg/kg LPS for eight weeks. During the modeling period, we observed the whole body weight of the HFD+LPD mice is gradually increased with the significant difference from the AL mice (Fig. 1a). After daily intraperitoneal injection with 0.25 mg/kg AS or 6 mg/kg NG for two weeks, it was seen that the HFD+LPS mice show a significant increase in the ratio of adipose tissue weight to the whole body weight (adipose weight/body weight ratio), whereas AS/NG allows a remarkable decrease in the adipose weight/body weight ratio in the HFD+LPS+AS mice (decrease for 37.5%) and HFD+LPS+NG mice (decrease for 25%). On the other hand, it was also noticed that the ratio of hepatic tissue weight to the whole body weight (hepatic weight/body weight ratio) is almost unchanged in the HFD+LPS+AS and HFD+LPS+NG mice (Fig. 1b).

**Figure 1.**
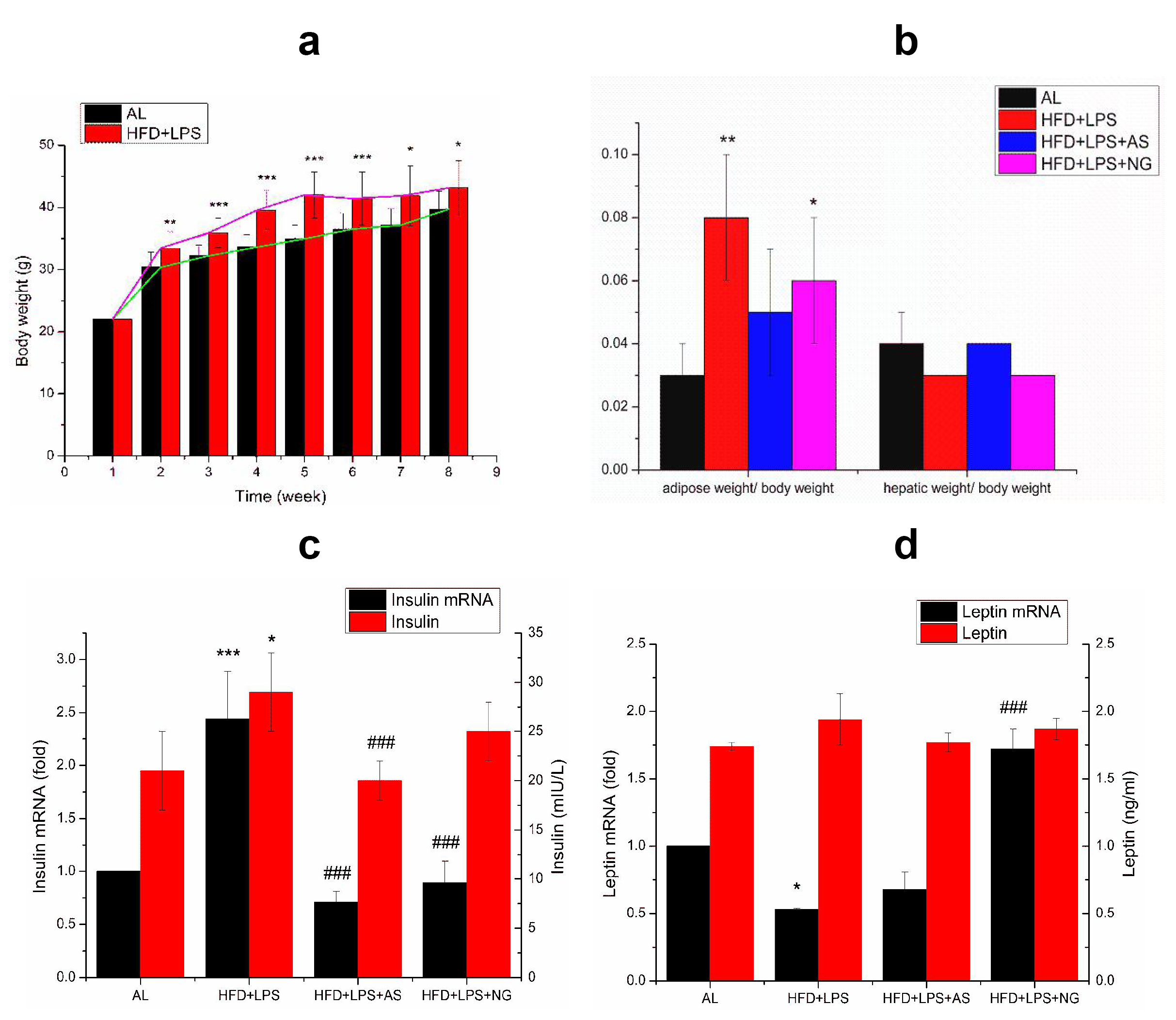
Comparison of AS/NG-mediated adipose weight reduction with low insulin and high leptin levels among AL, HFD+LPS, HFD+LPS+AS/NG mice. (**a**) The whole body weight curve. (**b**) The adipose weight/body weight ratio and the hepatic weight/body weight ratio. (**c**) The pancreatic insulin mRNAand serum insulin levels. (**d**) The adipose leptin mRNAand serum leptin levels.*** P<0.001 with very very significant difference from AL;** P<0.01 with very significant difference from AL;* P<0.05 with significant difference from AL;^###^ P<0.001 with very very significant difference from HFD+LPS;^##^ P<0.01 with very significant difference from HFD+LPS; # P<0.05 with significant difference from HFD+LPS.

We further quantified the pancreatic insulin mRNA levels and parallelly detected the serum insulin levels among all tested mice. Consequently, insulin mRNA levels are elevated in the HFD+LPS mice, but declined in the HFD+LPS+AS and HFD+LPS+NG mice. Accordingly, insulin levels are also elevated in the HFD+LPS mice, but declined in the HFD+LPS+AS and HFD+LPS+NG mice (Fig. 1c). Upon quantification of the adipose leptin mRNA levels and parallelly detection of the serum leptin levels, it was observed that the leptin mRNA levels and the leptin levels are almost unchanged among all tested mice (Fig. 1d). It could be concluded that AS/NG modulates a stable circulation of the insulin levels, which should ensure the metabolic homeostasis upon a conversion from insulin tolerance to insulin sensitivity.

### AS/NG lowers NIDDM and NAFLD risks and compromises inflammatory hepatic lesions

To evaluate whether AS/NG-mediated weight reduction effects would decrease the risk of fatty liver and type 2 diabetes, we quantitatively analyzed the expression profiles of 84 NAFLD/NIDDM-related genes using a tailored PCR microarray. Consequently, AS/NG was found to significantly downregulate NAFLD/NIDDM-related genes in the hepatic tissue (Fig. 2a and 2b). The AS/NG-treated HFD+LPS mice show more blue boxes representing the decreased NAFLD/NIDDM-related mRNAs than the HFD+LPS mice, whereas the HFD+LPS mice display more yellow box representing the increased NAFLD/NIDDM-related mRNAs than the AL mice.

**Figure 2.**
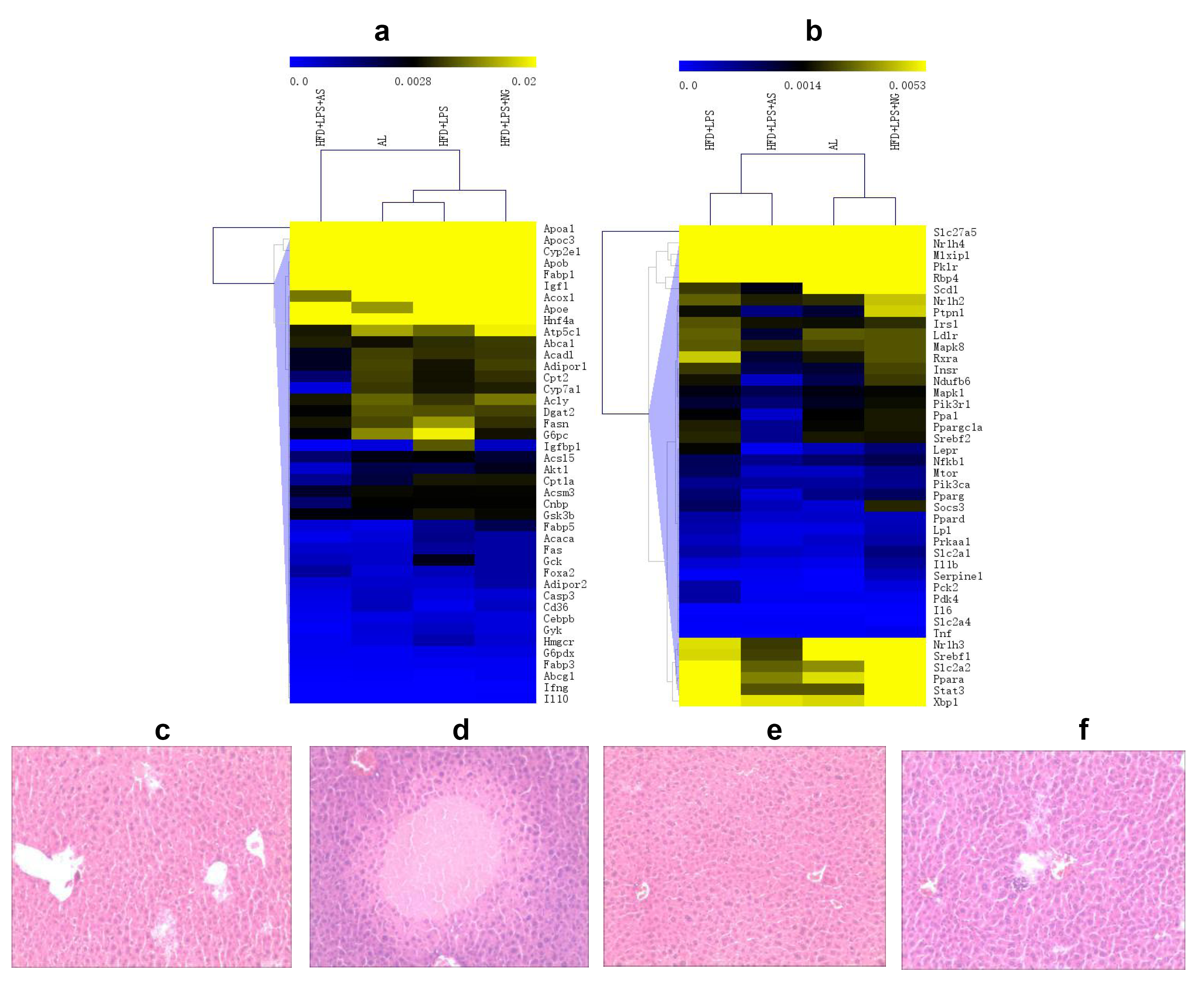
Comparison ofAS/NG-mediated lowered NIDDM/NALFD risks and less pathological alterations among AL, HFD+LPS, HFD+LPS+AS/NG mice. (**a**) The hepatic NIDDM/NAFLD-related mRNA levels (part 1). (**b**) The hepatic NIDDM/NAFLD-related mRNA levels (part 2). (**c-f**) HA-stained hepatic tissues fromAL, HFD+LPS, HFD+LPS+AS and HFD+LPS+NG mice (10×10).

The magnitudes of HFD+LPS-upregulated NIDDM-related genes were listed in Table 1. From the fold changes of mRNA levels, it was unambiguously seen that most NIDDM-related genes are downregulated, leading to their mRNA levels are lower than those in the HFD+LPS mice. For example, the mRNA levels of *Insr, Irs1, Mtor, Pik3ca(p110A),* and *Pik3r1(pi3k p85a),* whose encoding proteins belonging to the PI3K-Akt-mTORC1 signaling pathway, are higher in the HFD+LPS mice than in the HFD+LPS+AS mice or partly in the HFD+LPS+NG mice. These results demonstrated that AS/NG has decreased the risk of fatty liver and type 2 diabetes, at least on the mRNA levels.

As compared with the AL mice (Fig. 2c), a hepatic sample from the HFD+LPS mice exhibits significant focal necrosis, accounting for five scores (Fig. 2d), whereas the hepatic samples from AS/NG-treated HFD+LPS mice exhibit improvement without notable inflammatory filtration (Fig. 2e and 2f). These results indicated AS/NG treatment allows HFD+LPS mice recovery from the inflammatory hepatic lesions.

### AS/NG compromises NF-κB/iNOS signaling and downregulates pro-inflammatory cytokines

To make clear whether the visceral weight gain and visceral weight loss might be correlated with the pro-inflammatory and anti-inflammatory responses, we compared the expression levels of NF-κB in the hepatic tissues among all tested mice. As expectation, the HFD+LPS mice exhibit a higher *NF-κB* mRNA level than the AL mice, whereas the AS/NG-treated HFD+LPS mice show a lower *NF-κB* mRNA level than the HFD+LPS mice. In similar, the HFD+LPS mice possess a higher NF-κB level than the AL mice, whereas the AS/NG-treated HFD+LPS mice display an equal or lower NF-κB level than the HFD+LPS mice (Fig. 3a). Considering NF-κB induces *NOS2,* we anticipated AS/NG should downregulate *NOS2.* Indeed, it was observed that the hepatic *NOS2* mRNA and iNOS levels are higher in the HFD+LPS mice than in the AL mice, whereas the hepatic *NOS2* mRNA and iNOS levels become much lower after treatment of HFD+LPS mice by AS/NG (Fig. 3b).

**Figure 3.**
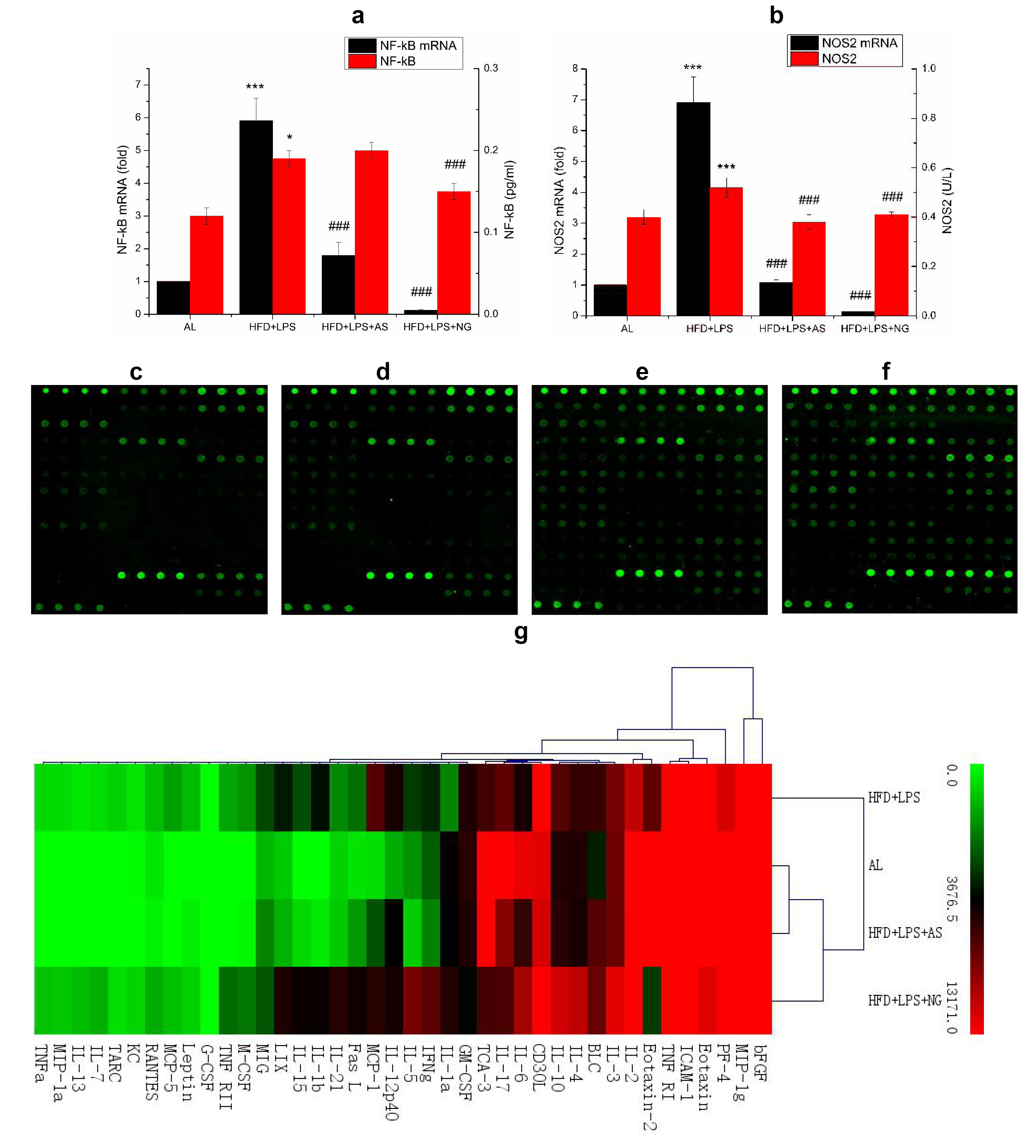
Comparison of AS/NG-induced *NF-κB* mRNA/NF-κB levels and pro-inflammation cytokine mRNA levels among AL, HFD+LPS, HFD+LPS+AS/NG-treated mice. (**a**) The hepatic *NF-κB* mRNA and NF-κB levels; (**b**) The hepatic *NOS2* mRNA and iNOS levels; (**c-f**) The PCR-based microarrays of the selective cytokine/chemokine and corresponding receptor mRNAs in the adipose tissues of AL, HFD+LPS, HfD+LPS+AS, and HFD+LPS+nG mice; (g) Quantification ofthe selective cytokine/chemokine and corresponding receptor mRNAs in the adipose tissues of AL, HFD+LPS, HFD+LPS+AS, and HFD+LPS+NG mice.*** P<0.001 with very very significant difference from AL;** P<0.01 with very significant difference from AL;* P<0.05 with significant difference from AL;^###^ P<0.001 with very very significant difference from HFD+LPS;^##^ P<0.01 with very significant difference from HFD+lPs;^#^ P<0.05 with significant difference from HFD+LPS.

**Table 1.**
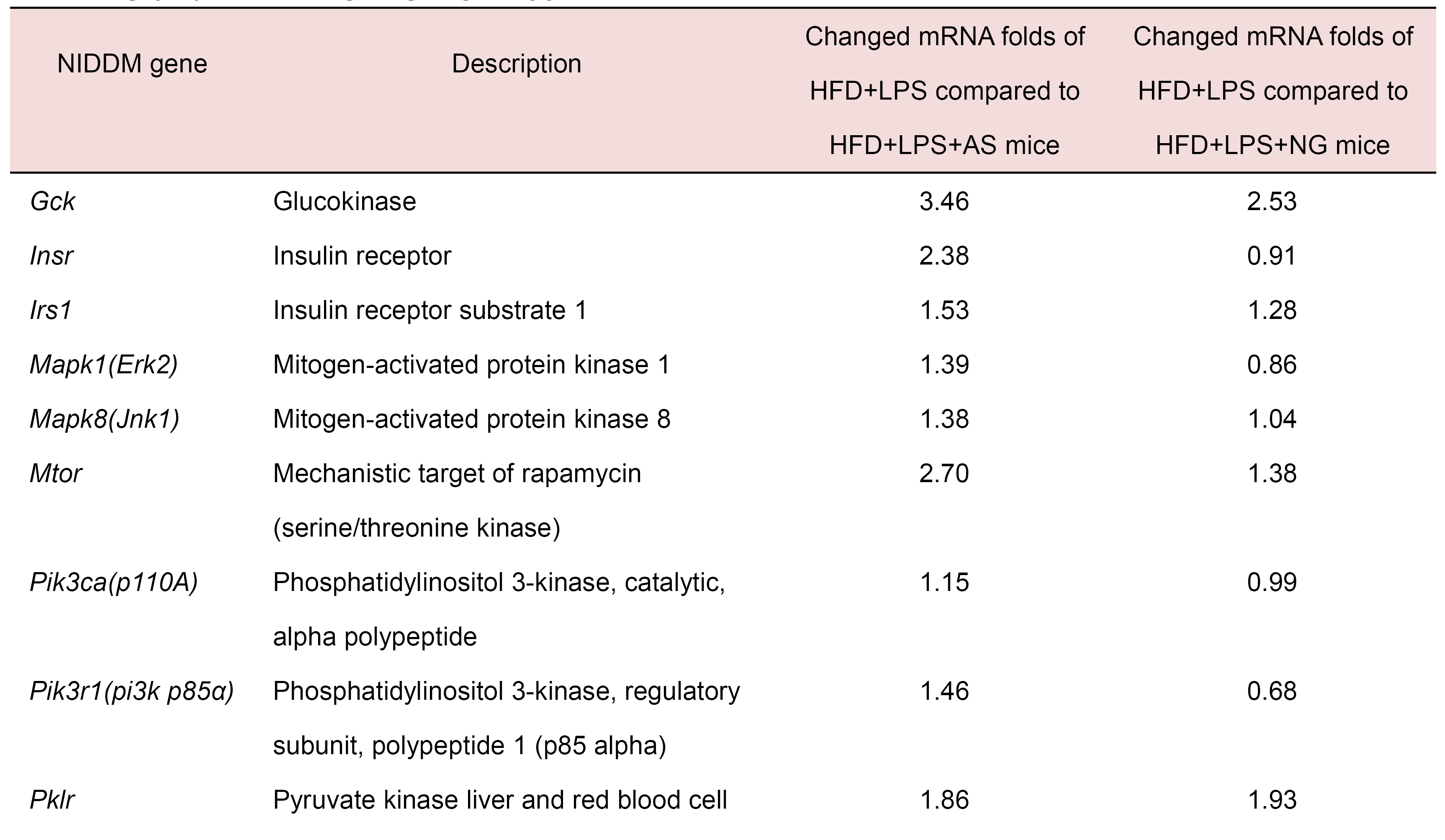
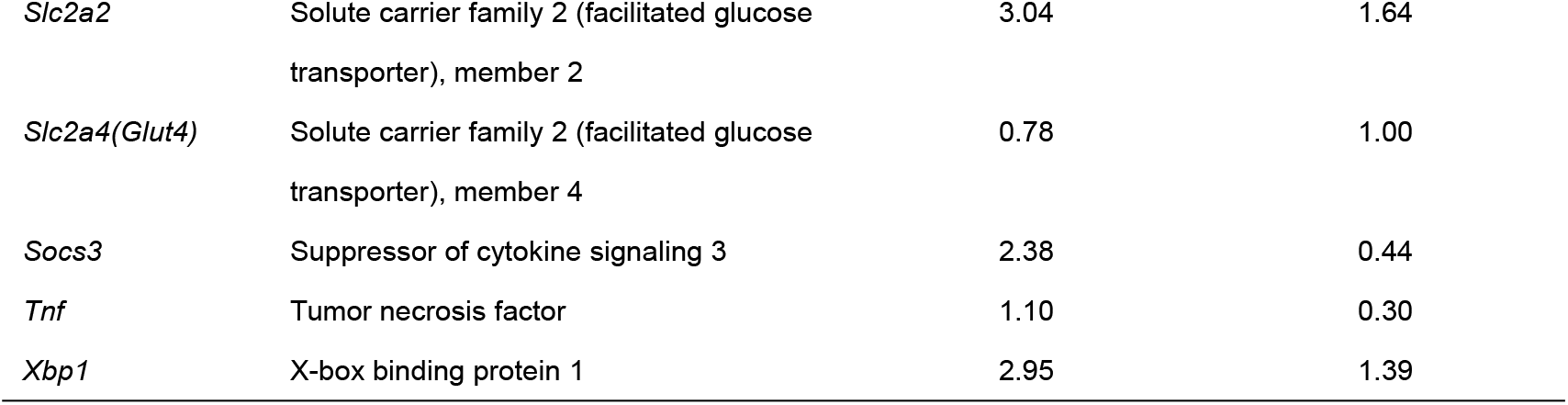
Effects of HFD+LPS and AS/NG on the hepatic NIDDM-related mRNA levels among HFD+LPS and HFD+LPS+AS/NG mice.

To figure out whether AS/NG would exert an anti-inflammatory effect in the visceral organs, we monitored the global changes of 200 cytokines/chemokines in the adipose tissues among all tested mice. As results, AS/NG tremendously affects the expression profiles among the shown cytokines/chemokines (see Fig. S1 for others), especially downregulates pro-inflammatory cytokines (Table 2). The AL mice (Fig. 3c) show an almost similar expression profile with the AS-treated HFD+LPS mice (Fig. 3d), whereas the HFD+LPS mice (Fig. 3e) exhibit a nearly identical expression pattern to the NG-treated HFD+LPS mice (Fig. 3f). In details, as shown in Fig. 3g, the pro-inflammatory cytokine *TNF-α* mRNA levels were undetectable in the AL and HFD+LPS+AS samples, but elevated for 546 folds in the HFD+LPS sample and 891 folds in the HFD+LPS+NG sample. The other pro-inflammatory cytokine *IL-1β* mRNA levels were undetectable in the AL sample, but elevated for 296 folds, 3477 folds, and 3914 folds in HFD+LPS+AS, HFD+LPS, and HFD+LPS+NG samples, respectively. On the other hand, from a view of the anti-inflammatory cytokine IL-10, its mRNA was accounted to an increase for 11435 folds in the HFD+LPS+NG sample, as compared to an increase for 4921 folds in the AL sample, 5267 folds in the HFD+LPS+AS sample, and 6867 folds in the HFD+LPS sample.

**Table 2:** A square matrix ofthe selected cytokine/chemokine and corresponding receptor mRNAs in the adipose tissues of AL, HFD+LPS, HFD+LPS+AS, and HFD+LPS+NG mice.

|  | 1,2,3,4 | 5,6,7,8 | 9,10,11,12 |
| --- | --- | --- | --- |
| a | POS1 | POS2 | bFGF |
| b | BLC | CD30L | Eotaxin |
| c | Eotaxin-2 | Fas L | G-CSF |
| d | GM-CSF | ICAM-1 | IFNγ |
| e | IL-1a | IL-1b | IL-2 |
| f | IL-3 | IL-4 | IL-5 |
| g | IL-6 | IL-7 | IL-10 |
| h | IL-12p40 | IL-13 | IL-15 |
| i | IL-17 | IL-21 | KC |
| j | Leptin | LIX | MCP-1 |
| k | MCP-5 | M-CSF | MIG |
| l | MIP-1a | MIP-1g | PF-4 |
| m | RANTES | TARC | TCA-3 |
| n | TNF RI | TNF RII | TNFα |

These results suggested that AS might exert the anti-inflammatory effects by downregulation of the pro-inflammatory cytokines including TNF-α and IL-1β, whereas NG might play the anti-inflammatory roles by upregulation of the anti-inflammatory cytokines such as IL-10.

### AS/NG modulates the expression of *NOS3/eNOS,* angiogenesis-related genes, and oxidative stress-responsive genes

To investigate whether downregulation of *NOS2/iNOS* might upregulate NOS3/eNOS, we detected the hepatic *NOS2* mRNA/iNOS levels in all tested mice. Indeed, a higher *NOS3* mRNA level (Fig. 4a) and a higher eNOS level (Fig. 4b) were measured in the HFD+LPS+AS mice. However, only an elevated eNOS level was determined in the HFD+LPS+NG mice. Surprisingly, a high hepatic eNOS level was also examined in the HFD+LPS mice. These results unnecessarily provided evidence supporting a causal relationship between NOS2/iNOS downregulation and NOS3/eNOS upregulation, suggesting an alternative modulation of iNOS and eNOS, such as changing their relative activities via the availability of L-arginine, a precursor for NO biosynthesis.

**Figure 4.**
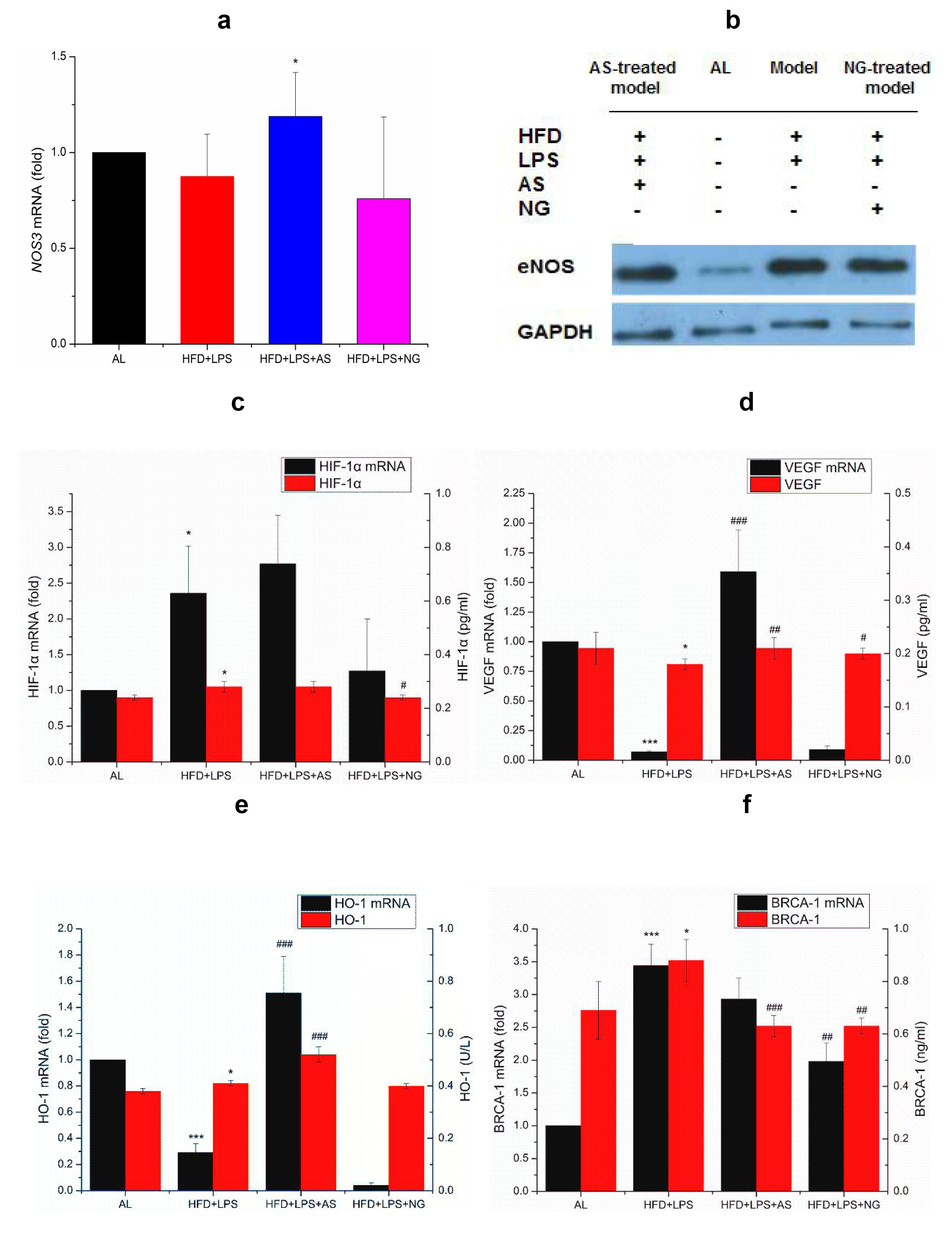
Comparison of AS/NG-induced angiogenesis-related genes and oxidative stress-responsible genes among AL, HFD+LPS, HFD+LPS+AS/NG-treated mice. (**a**) The hepatic *NOS3* mRNA level. (**b**) The hepatic eNOS level. (**c**) The hepatic *HIF-1α* mRNA and HIF-1α levels; (**d**) The hepatic *VEGF* mRNA and VEGF levels; (**e**) The hepatic *HO-1* mRNA and HO-1 levels; (**f**) The hepatic *BRCA1* mRNA and BRCA1 levels.*** P<0.001 with very very significant difference from AL; ** P<0.01 with very significant difference fromAL;* P<0.05 with significant difference fromAL;^###^ P<0.001 with very very significant difference from HFD+LPS;^##^ P<0.01 with very significant difference from HFD+LPS;^#^ P<0.05 with significant difference from HFD+LPS.

Owing to downregulation of iNOS and mitigation of NO burst, AS/NG was believed to allow O_2_ entry into COX, thereby attenuating hypoxia and angiogenesis. As expectation, AS/NG declines the hepatic HIF-1α levels (Fig. 4c) and VEGF levels (Fig. 4d) although their corresponding mRNA levels are still elevated. Accordingly, HO-1 that degrades the pro-oxidative heme (Fig. 4e) and BRCA1 that repairs oxidative DNA damage (Fig. 4f) are also downregulated by AS/NG, suggesting an essential outcome of mitigated ROS burst and compromised DNA damage.

### AS/NG induces mitochondrial biomarker expression, triggers NO generation, and enhances ATP production

Given AS mimics NG to rectify the mitochondrial dysfunction, we assumed AS/NG would drive mitochondrial biogenesis. To confirm this assumption, we intramuscularly injected mice with AS (0.25 mg/kg) and quantified the expression profiles of three mitochondrial structural and functional biomarkers, manganese SOD (Mn-SOD), cytochrome *c* oxidase 4 (COX4), and uncoupling protein 1 (UCP1) in mouse muscular cells. As depicted in Fig. 5a, it was noted that AS/NG significantly increases the muscular amounts of Mn-SOD, COX4, and UCP1. Furthermore, the non-specific NOS inhibitor N^G^-monomethyl-L-arginine monoacetate (L-NMMA) was found to repress AS/NG-induced upregulation of Mn-SOD, COX4, and UCP1, implying NO might be involved in mitochondrial biogenesis.

**Figure 5.**
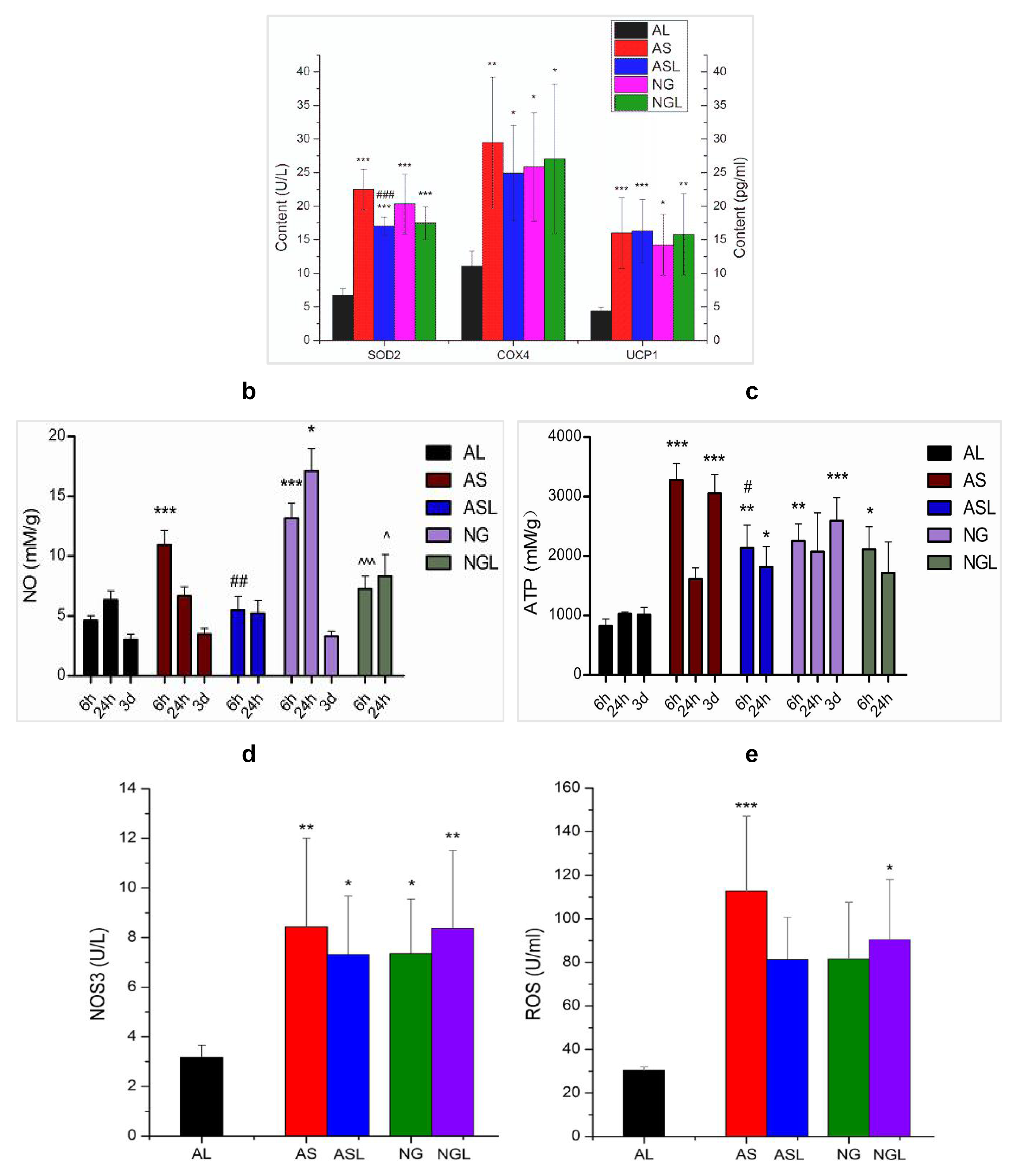
Comparison of AS/NG-induced upregulation of mitochondrial biomarkers accompanying with NO generation, ATP production, NOS3/eNOS upregulation, and ROS burst in AL, AS, ASL, NG, and NGL mice. (**a**) The muscular Mn-SOD, COX4, and UCP1 levels. (**b**) The muscular NO levels. (c) The muscular ATP levels. (**d**) The muscular NOS3/eNOS levels. (**e**) The muscular ROS levels. Samples were collected from mouse skeletal muscles after 6 hours by one injection or by daily injections for 3 days by 0.25 mg/kg AS or 6 mg/kg NG with or without 3 mg/kg L-NMMA. ASL designates AS supplemented with L-NMMA, and NGL designates NG supplemented with L-NMMA.*** P<0.001 with very very significant difference from AL;** P<0.01 with very significant difference from AL;* P<0.05 with significant difference from AL;^###^ P<0.001 with very very significant difference from AS in ASL;^##^ P<0.01 with very significant difference from AS in ASL;^#^ P<0.05 with significant difference from AS in ASL; ^ˆˆˆ^ P<0.001 with very very significant difference from NG in NGL. ^ˆˆ^ P<0.01 with very significant difference from NG in NGL; ^ˆ^ P<0.05 with very significant difference from NG in NGL.

**Figure 5.**
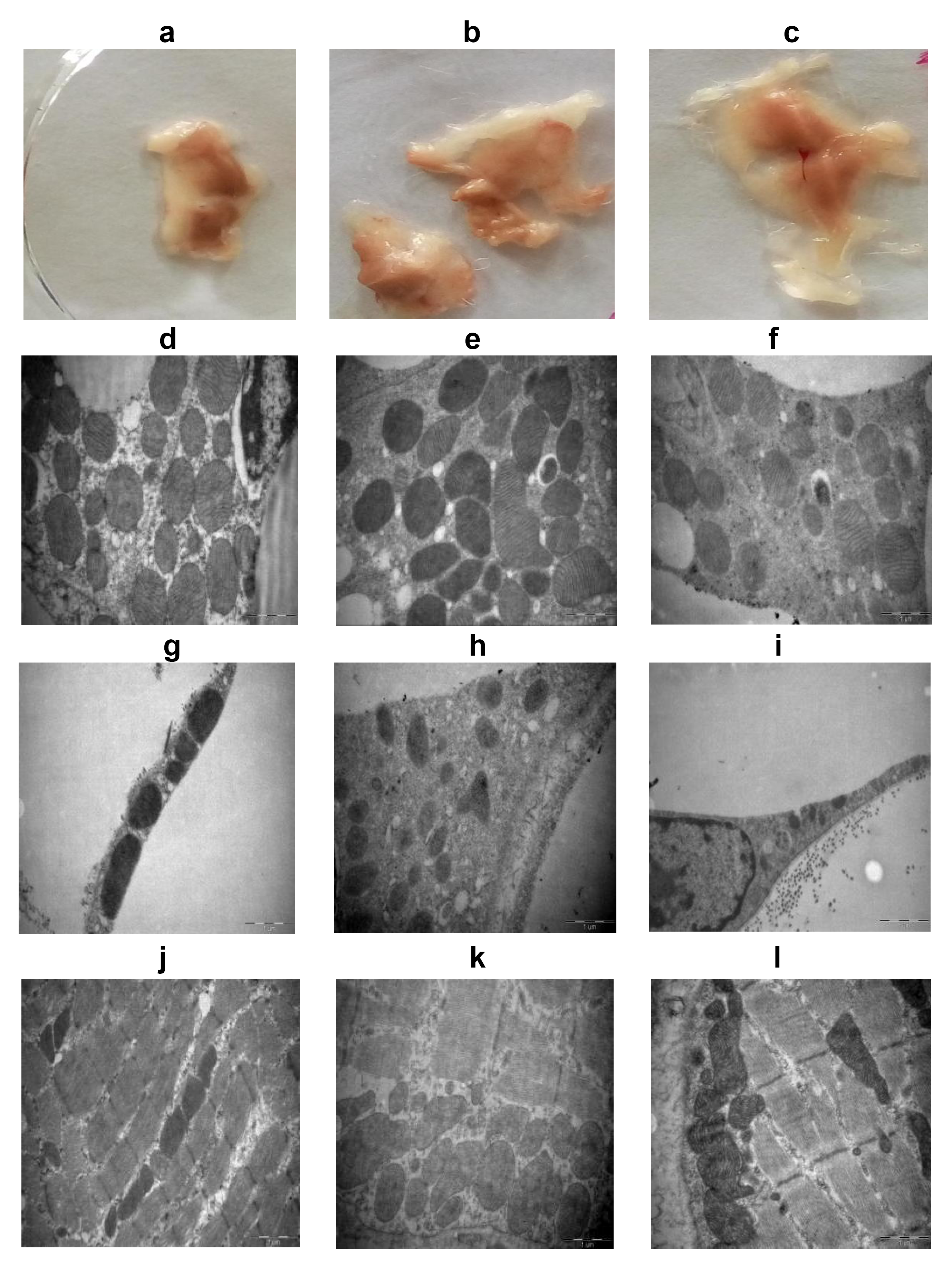
Comparison of AS/NG-triggered mitochondrial multiplying and adipose browning among AL, AS, and NG mice. (**a-c**) The morphological presentation of BAT. (**d-f**) The microscopic photo of mitochondria in BAT cells (50,000^×^). (**g-i**) The microscopic photo of mitochondria in WAT cells (50,000^×^). (**J-l**) The microscopic photo of mitochondria in the muscular cells (50,000^×^).

It was really noticed that the muscular NO levels are significantly increased upon AS/NG treatment in a time-dependent manner, but ASL/NGL generates the relatively lower NO levels (Fig. 5b). Given AS/NG also improves mitochondrial functions, we anticipated AS/NG should promote ATP production. Indeed, we were aware that while AS/NG gives rise to the higher ATP levels, AL and ASL/NGL generate the lower ATP levels (Fig. 5c), strengthening AS might mimic NG-released NO to impede mitochondrial respiration at first, but secondly to recover mitochondrial respiration. To chase the source of NO, we further determined the expression levels of NOS3/eNOS. As illustrated in Fig. 5d, AS/NG dramatically increases the muscular amount of eNOS whenever L-NMMA was supplemented or not. Accordingly, AS/NG and ASL/NGL can elicit ROS generation synchronous to eNOS induction (Fig. 5e), highlighting AS/NG transiently interrupts electron transport to trigger ROS burst.

### AS/NG triggers mitochondrial biogenesis and driven adipose conversion

To morphologically reveal whether AS/NG would trigger mitochondrial biogenesis, we investigated the phenotypical changes of mitochondria in the brown adipose tissue (BAT), white adipose tissue (WAT), and muscular tissue of the AS/NG-treated mice. As results, we saw the size of BAT from an AL mouse (Fig. 6a) is small than that from an AS mouse (Fig. 6b) or an NG mouse (Fig. 6c). In BAT, AL cells (Fig. 6d) show less dense (deep color) mitochondria than AS/NG cells (Fig. 6e), and the copy numbers of mitochondria in AS/NG cells (Fig. 6f) seem to be slightly higher than those in AL cells. In WAT, AL cells (Fig. 6g) possess only linear mitochondria, but AS cells (Fig. 6h) and NG cells (Fig. 6i) exhibit an expanded or enlarged mitochondrial area. As compared with the one-layer array of mitochondria in AL muscular cells (Fig. 6a), AS cells (Fig. 6b) and NG cells (Fig. 6c) exhibit the multi-layer array of mitochondria, suggesting a remarkable mitochondrial proliferation. These results indicated that AS resembles NG to trigger mitochondrial biogenesis, during which AS seems more effective than NG in promoting mitochondrial biogenesis.

**Figure 6.**
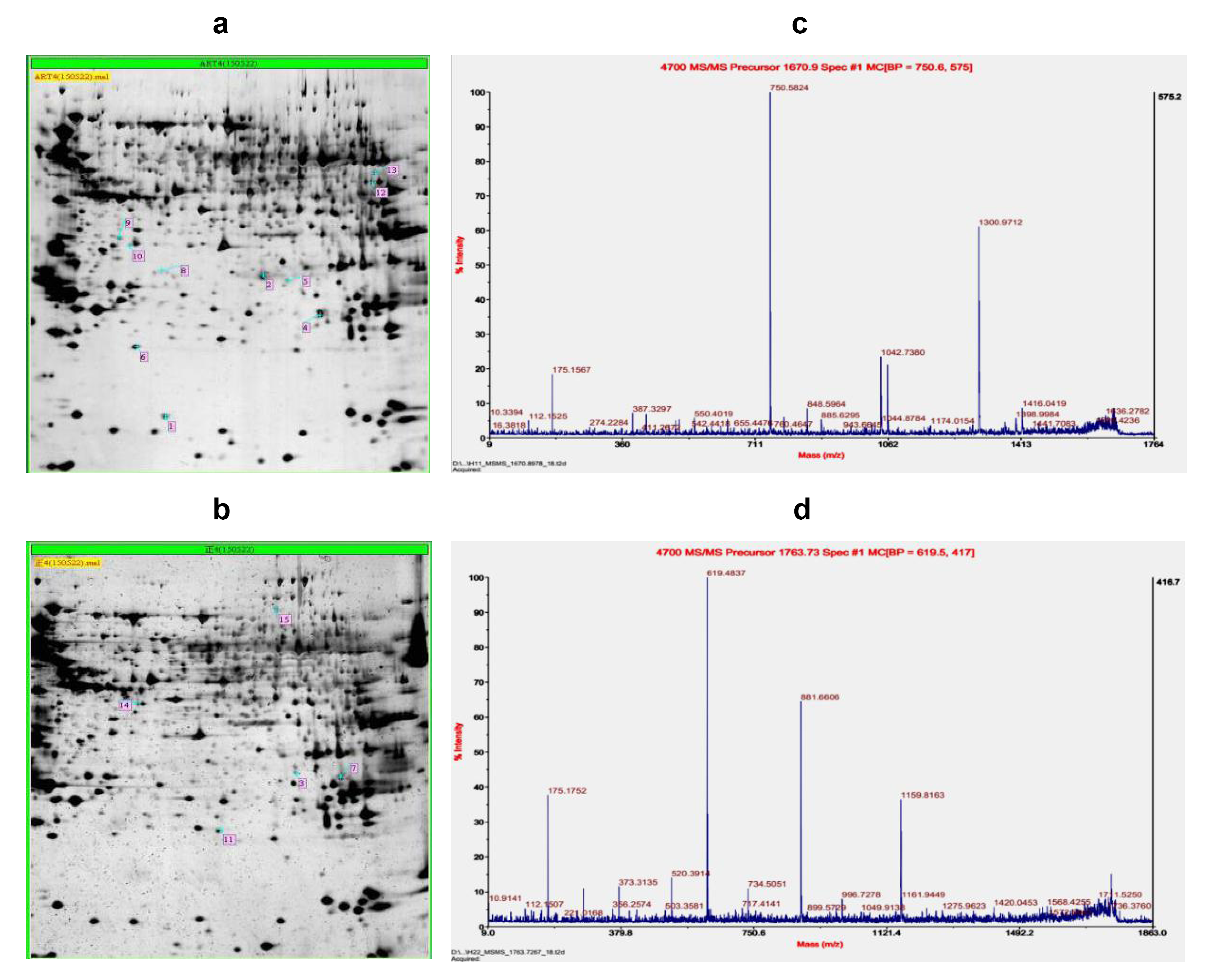
Identification of AS-targeted mitochondrial CYC1 and NDUFV1 in mouse pancreatic tissues by 2DE-MALDI-TOF-MS analysis. (**a**) The 2DE gel image of pancreatic proteins separated from an AS-treated mouse sample. (**b**) The 2DE gel image of pancreatic proteins separated from an AL mouse sample. (**c**) The peptide map of CYC1 (spot2/H11). (**d**) The peptide map of NDUFV1 (spot 13/H22).

### AS targets mitochondrial CYC1 and NDUFV1

To identify AS-targeted cytosolic and mitochondrial proteins, we choose pancreatic tissues for protein extraction and identification after intraperitoneal injection of mice with 0.25 mg/kg AS for three days. Total proteins from an AS sample were separated by two-dimensional electrophoresis (2DE), from which 10 differential proteins with the sample number/spot number, 1/H10, 2/H11, 4/H13, 5/H14, 6/H15, 8/H17, 9/H18, 10/H19, 12/H21, and 13/H22, were labeled on the gel (Fig. 7a). Parallelly, five protein spots from an AL sample, 3/H12, 7/H16, 11/H20, 14/H23, and 15/H24, were also identified after running the total proteins on the gel (Fig. 7b). Clearly, those 15 differential protein spots should represent upregulated or downregulated proteins.

For further identification, all those 15 protein spots were excised from the gels for analysis by the matrix assisted laser desorption ionization time of flight mass spectrometry (MALDI-TOF-MS). Consequently, 12 protein spots (1/H10, 2/H11, 3/H12, 4/H13, 6/H15, 7/H16, 9/H18, 11/H20, 12/H21, 13/H22, 14/H23, and 15/H24) were successfully identified (see Supplementary Table 2). While two protein spots, 2/H11 and 13/H22, were verified to be the mitochondrial proteins with unique peptide maps, CYC1 and NDUFV1, other 10 protein spots were classified as cytosolic proteins with multiple peptide maps. Importantly, only glutathione S transferase (4/H13) and pancreatic triacylglyceral lipase (15/H24) are among the candidate AS-binding targets with the highest protein scores (see Fig. S2). The detailed peptide maps of AS-targeted CYC1 and NDUFV1 were shown in Fig. 7c and 7d, and their molecular indice were summarized in Table 3.

**Table 3:** Identification of AS-targeted mitochondrial CYC1 and NDUFV1 in mouse pancreatic tissues.

| Gel Index/<br>Position | Rank | Protein Name | Accession<br>No. | Protein<br>ID | Protein<br>MW | Protein<br>Score | Protein<br>Score C.I.% |
| --- | --- | --- | --- | --- | --- | --- | --- |
| 184/H11 | 1 | Unnamed protein product (CYC1) | gi\|74213069 | BAE41677.1 | 29478.7 | 94 | 99.994 |
|  | 2 | CYC1 | gi\|52350626 | AAH82790.1 | 34383.1 | 91 | 99.988 |
|  | 3 | Unnamed protein product (CYC1) | gi\|12833077 | BAB22380.1 | 35418.0 | 90 | 99.987 |
| 195/H22 | 1 | NDUFV1 isoform CRA_a | gi\|148701042 | EDL32989.1 | 36315.1 | 155 | 100 |
|  | 2 | NDUFV1 mitochondrial precursor | gi\|19526814 | NP598427.1 | 50801.7 | 147 | 100 |

C.I.: confidence identification; CPD: conserved protein domain.

These results demonstrated AS targets the mitochondrial proteins in addition to the cytosolic proteins, addressing a mechanism underlying AS-mimicking NO to elicit mitochondrial biogenesis and convert adipose energy deposition to adipose energy expenditure.

## Discussion

In accordance with the practice from other authors that infiltration by inflammatory cells into the hepatic tissues of HFD-fed mice occurs after treatment of mice by 0.25 mg/kg LPS^26^, our previous work also unraveled HFD-combined low-dose (0.25 mg/kg) LPS induces obesity with a global upregulation of pro-inflammatory cytokines, whereas HFD alone or HFD-combined high-dose (1.2 mg/kg) LPS only induces obesity without an extensive upregulation of pro-inflammatory cytokines^27^. Upon HFD+0.25 mg/kg LPS treatment in the present study, we successfully established an inflammatory obese mouse model, in which the adipose weight/body weight ratios were increased, insulin levels elevated, and pro-inflammatory cytokines and NIDDM/NALFD-related genes upregulated. By daily administering the model mice with 0.25 mg/kg AS or 6 mg/kg NG for two weeks, we not only observed the reduced adipose weight/body weight ratios and declined insulin levels, but also noticed the downregulated pro-inflammatory cytokines and NIDDM/NALFD-related genes. As our knowledge, this is the first report describing an effect of artemisinin on adipose weight reduction although a potential of artemisinin in amelioration of the type 1 diabetes has been also shown in a distinct mechanism, by which artemisinin targets gephyrin to enhance GABAA receptor signaling and induce the conversion of pancreatic islet α-cells to insulin-secreting β-cells^28^.

Based on the previous reports indicating AS inhibits iNOS and NF-κB^29,30^, and from the present results revealing HFD+LPS-induced pro-inflammation prompts adipose weight gain and AS/NG-mediated anti-inflammation accelerates adipose weight loss, we have experimentally confirmed our suggested hypothesis that pro-inflammation/anti-inflammation-switched dysfunctional/functional mitochondria modulate energy deposition/expenditure. A conversion from dysfunctional mitochondria to functional mitochondria might be mediated by a shift from iNOS activated by pro-inflammation and pro-oxidation to eNOS activated by anti-inflammation and anti-oxidation^31^ because the former produced high-level NO can completely block oxidative phosphorylation, whereas the latter generated low-level NO can only partly interrupt electron transport. Logically, it should be that iNOS-derived high-level NO promotes energy deposition and weight gain, whereas eNOS-derived low-level NO enhances energy expenditure and weight loss. It was demonstrated by other authors that a chronic blockade of iNOS by L-NMMA can reduce adiposity and compromise insulin resistance in HFD-induced obese mice^32^. We also noticed AS/NG-driven *NOS3*/eNOS upregulation is correlated with ATP overproduction in the present study.

We surprisingly observed an elevated eNOS level in the HFD+LPS mice, implying *NOS3* is also upregulated under the pro-inflammatory condition. This phenomenon was also seen after use of the non-steroidal anti-inflammatory drugs (NSAIDs), in which oxidative stress leads to increase in eNOS expression but decrease in NO production^33^. It was deciphered by the so-called eNOS uncoupling, in which eNOS produces superoxide anion (O_2_^-^) instead of NO^34^. In the uncoupled eNOS, depletion of the co-factor tetrahydrobiopterin (BH_4_) may occur upon exposure to excessive peroxynitrite (ONOO^-^), hence leading to eNOS dysfunction^35^. On the other hand, eNOS can be modified post-translationally^36^, in which NO-mediated S-nitrosylation may inactivate eNOS and in turn upregulate *NOS3*^37^. Additionally, eNOS activity is dependent on the availability of the NO precursor *L*-arginine^38^, implying iNOS that dispenses large amounts of *L*-arginine should tremendously decrease eNOS activity. pro-inflammation also leads to Therefore, it was believed that eNOS can exert the normal physiological functions only in the anti-inflammatory milieu. We actually noticed *NOS3* upregulation, NO elevation, and ATP overproduction after injecting AS/NG into mice without inflammation.

It was previously indicated that adipose tissue expansion is essentially initiated from adipocyte inflammation^39^, highlighting an association of inflammation with visceral obesity. In such a context, it can be predicted that anti-inflammatory agents should ameliorate some obesity-related syndromes. Indeed, salicylate, a degraded product of the non-steroidal anti-inflammatory drug aspirin, was shown to reduce circulating lipids and increase insulin sensitivity in obese rats^40^. Similarly, nitro-aspirin was also proven to possess a potential to treat NAFLD^41^. In the present work, we observed AS/NG-induced anti-inflammatory responses decrease the risk of NAFLD/NIDDM. Some encouraging findings that underlie a beneficial effect of anti-inflammation on obesity-related disorders include aspirin ameliorates the type 2 diabetes as an activator of AMPK^42^, DNP induces pAMPK and p38 MAPKfor improvement of the type 2 diabetes and fatty liver^43^, and metformin and salicylate activate AMPK to inhibit liver lipogenesis and increase insulin sensitivity^44^. Additionally, we previously verified AS as well as the NO donor sodium nitroprusside and the NO precursor *L*-arginine extend mouse and yeast lifespan by tremendously upregulating AMPK/pAMPK^24^,^25^.

The covalent conjugation of artemisinin with haem was first identified in 1990’s, when the artemisinin-haem adducts were identified by mass spectrometry^45^,^46^. Later, artemisinin was verified to alkylate haem *in vitro* via dimethyl ester formation and dematallation^47^. The prosthetic haem groups harbouring in the haem proteins were validated as the cellular targets of artemisinin in mice^48^ and malarial parasites^49^. Based on a correlation of structure with function, two cytosolic haem proteins, NOS and catalase, from bacteria and tumor cells, as well as the mitochondrial haem protein COX in yeast were clarified as the targets of artemisinin *in vivo*^24^,^25^. Furthermore, as many as 124 malarial non-haem proteins were identified to covalently bind to artemisinin in *P. falciparum*^50^. Additionally, sarco/endoplasmic reticulum Ca^2^+-ATPase (SERCA)/PfATP6^51^, translational controlled tumor protein (TPCP)^52^, and glutathione S transferase (GST)^53^ were independently classified as artemisinin-conjugated non-haem proteins in the malarial parasite.

Using AS-injected mice, we ascertained AS not only targets the mitochondrial proteins, CYC1 and NDUFV1, but also targets the cytosolic proteins, GST and triacylglyceral lipase (TAGL). While the previously identified non-haem proteins NDUFV1 in yeast^17^ and GST in malaria^53^ were confirmed as the AS-targeted proteins in mice, the haem protein CYC1 and the non-haem protein TAGL were also identified as AS-targeted proteins in this study. As an essential outcome of NO interacting with COX, mitochondrial biogenesis should be initiated^54^. In the present study, enhanced mitochondrial biogenesis and recovered mitochondrial coupling were also validated by monitoring *COX4, SOD2,* and *UCP1* upregulation, ATP overproduction, and mitochondria replication. As previously indicated that decreased adipose mitochondria usually occur in the obese persons^55^, we also observed the conversion from white adipose tissues (WAT) with scarce mitochondria to brown adipose tissues (BAT) with abundant mitochondria in mice.

Regarding why the antimalarial agent AS can also exert weight-reducing effects, our answer is AS behaves in a dose-dependent manner, in which a high dose of AS can be used for antimalaria, while a trace amount of AS is suitable for weight reduction. In the present study, we employed quite a low dose of AS (0.25 mg/kg) for weight reduction. In our previous researches, we used a similar low dose of AS (0.3 mg/kg) to ameliorate articular synovitis in mice^56,57^. An *in vivo* pharmacokinetical investigation in mice demonstrated that 6.7 mg/kg AS leads to a peak serum level of 0.82 μg/ml AS in mice and rabbits after administration for 4 h^58^. This dose of AS is 5000 times the half inhibitory concentration (IC50) of *in vitro* tested *P. berghei,* and also close to the clinical dose of AS applied to malarial patients.

Taken together, we disclosed for the first time that AS exerts adipose weight-reducing effects and decreases NAFLD/NIDDM risks via enhancing a shift from pro-inflammation-switched aberrant mitochondrial functions that confer adipose weight gain to anti-inflammation-switched recovered mitochondrial functions that endow adipose weight loss. Our findings should denote so-called healthy or subcutaneous adipose depots caused as nutrient intake is more than energy expenditure and unhealthy or visceral adipose depots originated from inflammation.

## Methods

### Animals and treatment procedures

The female SWISS mice that belong to an out-bred population from SWISS mice were housed on a 12-h light and 12-h dark cycle at 25°C, and fed with HFD (60% basic feed-stuff + 20% lard + 10% sucrose + 10% yolk) or *ad libitum* chow (AL). For HFD+LPS modeling, mice were firstly fed with AL for 2 w, and then fed with HFD for 6 w, during which 0.25 mg/kg LPS was injected into peritoneal from the 5^th^ week on every 2 d for 2 w. For drug treatment, mice were intraperitoneally injected daily with 0.25 mg/kg AS or 6 mg/kg NG for 2 w. Animal treatment procedures were in accordance with the animal care committee at the Guangzhou University of Chinese Medicine, Guangzhou, China. The protocol was approved by the Animal Care Welfare Committee of Guangzhou University of Chinese Medicine (Permit Number: SPF-2011007).

### Quantitative polymerase chain reaction (qPCR)

Total RNA was extracted by a Trizol method for subsequent reverse transcription (RT) and PCR reactions. A reference gene and each target gene were amplified with a pair of the following specifically designed primers. The copy numbers of amplified genes were estimated by 2^-ΔΔCt^, in which ΔΔCt = [target gene (treatment group) / target gene (control group)] / [reference gene (treatment group) / reference gene (control group)].

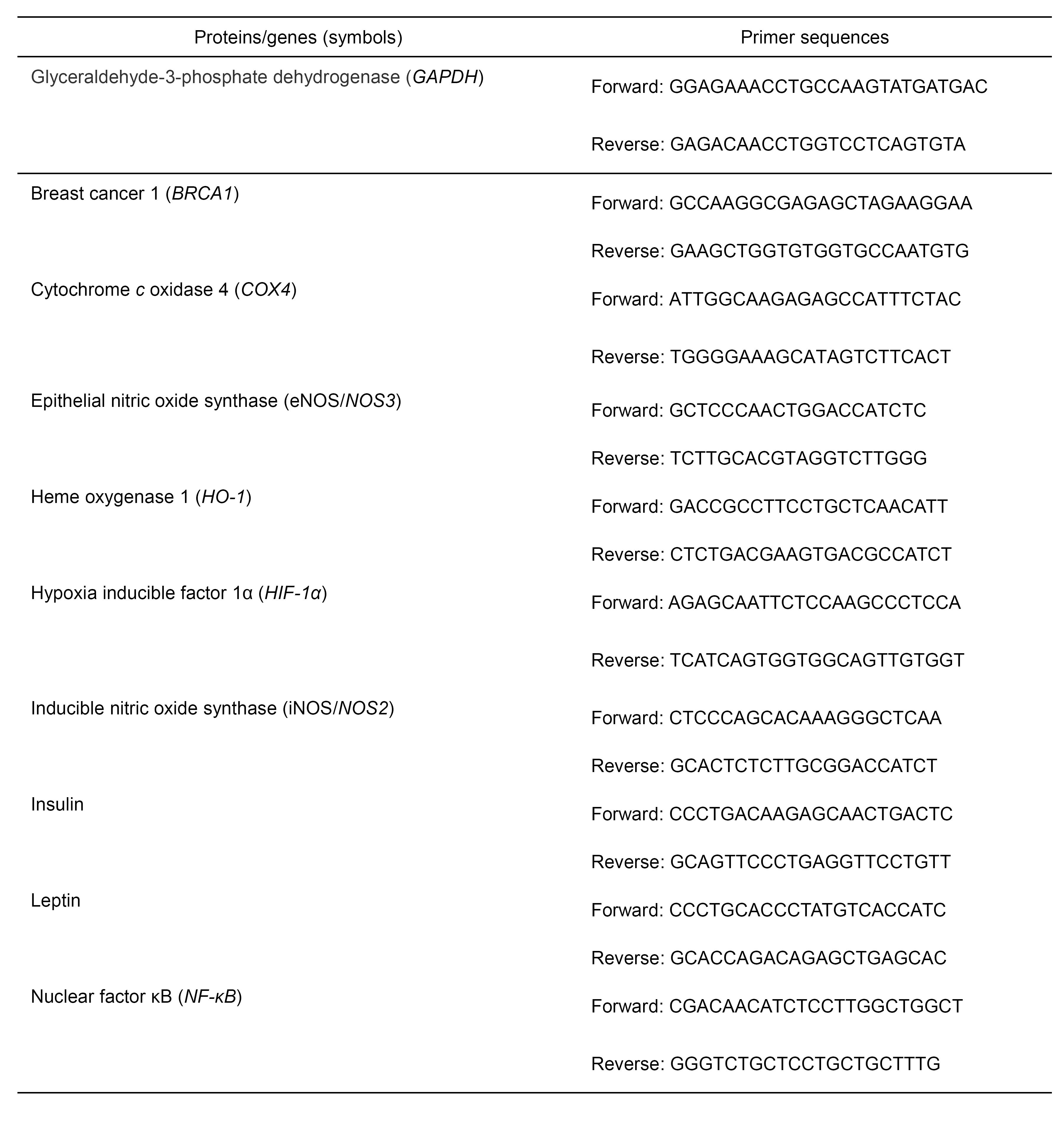

### PCR array and cytokine antibody array

The RT^2^ Profiler™ PCR Array Mouse Fatty Liver ( PAMM-157Z ) was purchased from SABioscience Qiagen, Hilden, Germany, and Quantibody^®^ Mouse Cytokine Antibody Array 4000 was purchased from RayBiotech, Inc, Norcross, GA, USA. The experiments were performed respectively by Kangchen Biotechnology Co, Ltd, Shanghai, China, and RayBiotech, Inc. Guangzhou, China. Protein extraction from blood cells by Cell & Tissue Protein Extraction Reagent (KangChen KC-415) was conducted according to the manufacturer’s instruction. Cytokine antibody array was carried out by Kangchen Bio-Tech, Shanghai, China using RayBio^®^ Mouse Cytokine Antibody Array.

### Enzyme-linked immunosorbent assay (ELISA) and Western blotting (WB)

All target proteins (insulin, leptin, NF-κB, iNOS, HIF-1α, VEGF, HO-1, BRCA1, SOD2, COX4, UCP1, eNOS) as compared to the reference protein GAPDH were immunoquantified by ELISA kits manufactured by Shanghai Yuanye Bio-Technology Co., Ltd, Shanghai, China. ROS levels were measured with a Mouse ROS ELISA Kit from EIAab Science Co. Ltd., Wuhan, China following the manufacturer’s instructions. The antibody against eNOS used for WB was purchased from Assay Biotechnology Co. Ltd., Sunnyvale, CA, USA.

### Electronic microscopy

After treatment, cells were harvested and fixed in 2.5% glutaraldehyde in 0.1 M phosphate buffer for three hours at 4 °C, followed by post-fixation in 1% osmium tetroxide for one hour. Samples were dehydrated in a graded series of ethanol baths, and infiltrated and embedded in Spurr’s low-viscosity medium. Ultra-thin sections of 60 nM were cut in a Leica microtome, double-stained with uranyl acetate and lead acetate, and examined in a Hitachi 7700 transmission electron microscope at an accelerating voltage of 60 kV.

### Two-dimensional electrophoresis (2DE)

The 2DE procedures including immobilized pH gradient (IPG) strip rehydration, the first dimensional isoelectric focusing (IEF), IPG strip equilibration, the second dimensional sodium dodecyl sulfate-polyacrylamide gel electrophoresis (SDS-PAGE), and silver staining/detection were mainly according to the instructions described in the Bio-Rad manual. Briefly, a total volume of 0.35 ml of suspension containing 0.8 mg total proteins was added into gel wells, and re-swelled for 12 h at 30 V. After the completion of the first dimensional IEF up to 80 kV ■ h, the IPG strip was immediately equilibrated for two times with 15 min each. The equilibrated strip was transferred to the protean II xi cell for the second dimensional SDS-PAGE. The concentration of polyacrylamide gel was 12%, and the condition of electrophoresis was first at 10 mA for 30 min and then at 25 mA until bromide blue reaches the gel bottom. Finally, the gel was stained by silver, scanned, and analyzed by the Image Master 2D Elite 5.0 software.

### Matrix-assisted laser absorption/ionization time of flight/time of flight mass spectrometry (MALDI-TOF/TOF MS)

For MALDI-TOF/TOF MS analysis, protein digestion extracts (tryptic peptides) were resuspended with 5 μl of 0.1 % trifluoroacetic acid and then the peptide samples were spotted onto stainless steel sample target plates and mixed (1:1 ratio) with a matrix consisting of a saturated solution of α-cyano-4-hydroxy-transcinnamic acid in 50% acetonitrile-1% trifluoroacetic acid. Peptide mass spectra were obtained on an Applied Biosystem Sciex 4800 MALDI TOF/TOF mass spectrometer. Data were acquired in positive MS reflector using a CalMix5 standard to calibrate the instrument (ABI4700 Calibration Mixture). Mass spectra were obtained from each sample spot by accumulation of 600 laser shots in an 800-4000 mass range. For MS/MS spectra, the 7 most abundant precursor ions per sample were selected for subsequent fragmentation, and 900-1200 laser shots were accumulated per precursor ion. The criterion for precursor selection was a minimum S/N of 50. For database searching and analysis, both the MS and MS/MS data were interpreted and processed by using the GPS Explorer software (V3.6, Applied Biosystems), then the obtained MS and MS/MS spectra per spot were combined and submitted to MASCOT search engine (V2.1, Matrix Science, London, U.K.). Mascot protein score with 95% Confident Identification (CI) was accepted.

### Histopathological analysis

A piece of the tissue was fixed by 10% formaldehyde followed by paraffin embedding and haematoxylin-eosin (HE) staining. The degenerative scores of inflammatory lesions were recorded as: 1-2 points indicate mild/severe hydropic degeneration; 3-4 points indicate mild/severe adipose degeneration; 5 points indicate necrosis.

### Spectrophotometry

The NO and ATP levels were determined using the reagent kits manufactured by Jiancheng Biotechnology Institute, Nanjing, China. All determination procedures were according to manufacturers instructions. The NO level (μM) = (OD _test_ − OD _blank_) / (OD _standard_ − OD _blank_) ⋅ standard nitrate concentration (20 μM). The ATP concentration (mM/g protein) = (OD _test_ − OD _control_) / (OD _standard_ − OD _blank_) ⋅ standard ATP concentration (50 μM) • dilution folds / protein content (g proteins/L).

### Statistical analysis

The software SPSS 22.0 was employed to analyze data, and the software GraphPad Prism 5.0 was employed to plot graphs. The Independent Simple Test was used to compare all groups, but the Kruskal-Wallis Test followed by Nemenyi test was used when the data distribution is skewed. All data were represented as mean ± SEM unless otherwise stated. The significance level *(p* value) was set at <0.05 (*), <0.01 (**), and <0.001 (***).

## Acknowledgements

This work was supported by the National Natural Science Foundation of China (No. 81273620 to Qing-Ping Zeng, No. 81673861 to Chang-Qing Li, and No. 81273817 and No. 81473740 to Qi Wang), and Guangdong Science and Technology Plan Project (No. 20150404042 to Qin Xu). We thank our colleagues in the Tropical Medicine Institute and Clinical Pharmacology Institute, Guangzhou University of Chinese Medicine, China. We also thank Kangchen Biotechnology Co, Shanghai, China for performance of RT-PCR microarray experiments.

## Author contributions

QPZ and QX designed the study. QG, JH, TL, YPC, and LLT carried out the experiments. XAH performed bioinformatics analysis. JDZ, CQL, QZ, QW, and QX participated in the interpretation of results. QPZ and QX wrote the manuscript with input from other authors. All authors read and approved the final manuscript.

## Additional information

**Supplementary Information** accompanies this paper at

